# Intranasal Delivery of Lithium Salt Suppresses Inflammatory Pyroptosis in the brain and Ameliorates Memory Loss and Depression-like Behavior in 5XFAD mice

**DOI:** 10.1101/2024.09.18.613794

**Authors:** Piplu Bhuiyan, Wenjia Zhang, Ge Liang, Bailin Jiang, Robert Vera, Rebecca Chae, Kyulee Kim, Lauren St. Louis, Ying Wang, Jia Liu, De-Maw Chuang, Huafeng Wei

**Affiliations:** Department of Anesthesiology and Critical Care, Perelman School of Medicine, University of Pennsylvania, Philadelphia, PA 19104, U.S.A.; Department of Anesthesiology, Shandong Provincial Hospital Affiliated to Shandong First Medical University, Jinan, Shandong, 250021, China; Department of Anesthesiology, Peking University People’s Hospital, Beijing, China; Department of Anesthesiology, The Affiliated Hospital of Qingdao University, Qingdao, Shandong, 26600, P. R. China; Scientist Emeritus, Intramural Research Program, National Institute of Mental Health, National Institutes of Health, Bethesda, MD 20892, USA

**Keywords:** Lithium, pharmacokinetics, Alzheimer’s disease, therapy, cognition, depression behavior, calcium, InsP_3_Rs, reactive oxygen species (ROS), oxidative stress, neurodegeneration, side effect, organ toxicity

## Abstract

**Background:** Alzheimer’s disease (AD) is a devastating neurodegenerative disease (AD) and has no treatment that can cure or halt the disease progression. This study explored the therapeutic potential of lithium salt dissolved in Ryanodex formulation vehicle (RFV) and delivered to the brain by intranasal application. We first compared lithium concentrations in the brain and blood of wild-type mice following intranasal or oral administration of lithium chloride (LiCl) dissolved in either RFV or water. The beneficial and side effects of intranasal versus oral LiCl in RFV in these mice were assessed and potential mechanisms underlying the efficacy of anti-inflammation and anti-pyroptosis in the brains were also investigated in both wild-type (WT) and 5XFAD Alzheimer’s Disease (AD) mice brains.

**Methods:** For the study of brain versus blood lithium concentrations, WT B6SJLF1/J mice at 2 months of age were treated with intranasal or oral LiCl (3 mmol/kg) dissolved in RFV or in water. Brain and blood lithium concentrations were measured at various times after drugs administration. Brain/blood lithium concentration ratios were then determined. For studying therapeutic efficacy versus side effects and their underlying mechanisms, 5XFAD and WT B6SJLF1/J mice were treated with intranasal LiCl (3 mmol/kg) daily, Monday to Friday each week, in RFV beginning at 2 or 9 months of age with a 12-week treatment duration. Animal behaviors were assessed for depression (tail suspension), cognition (fear conditioning and Y maze), olfaction (buried food test), and motor functions (rotarod) at the age of 5 and 12 months. Blood and brain tissue were harvested from these mice at 13 months. Blood biomarkers for the functions of thyroid (thyroid stimulating hormone, TSH) and kidney (creatinine) were measured using ELISA. Changes in protein expression levels of the endoplasmic reticulum Ca^2+^ release channels type 1 InsP_3_ receptors (InsP_3_R-1), malondialdehyde (MDA)-modified proteins and 4-hydroxy-2-nonenal (4-HNE), pyroptosis regulatory proteins (NLR family pyrin domain containing 3 (NLRP3), cleaved caspase-1, N-terminal of Gasdermin D (GSDMD)), cytotoxic (IL-1β, IL-18, IL-6, TNF-α) and cytoprotective (IL-10) cytokines and synapse proteins (PSD-95, synapsin-1) were determined using immunoblotting. Mouse body weights were monitored regularly.

**Results:** Compared to oral LiCl in RFV nanoparticles, intranasal treatment of WT mice with LiCl in RFV markedly decreased blood concentrations at the time frame of 30-120 minutes. The ratio of brain/blood lithium concentration after Intranasal lithium chloride in RFV significantly increased, in comparison to those after oral administration lithium chloride in RFV or intranasal administration of lithium chloride in water. Intranasal lithium chloride in RFV inhibited both memory loss and depressive behavior in adult and aged 5XFAD mice. Additionally intranasal treatment of aged 5XFAD mice with LiCl in RFV effectively suppressed the increases in InsP_3_R-1, intracellular oxidative stress markers (4-HNE-bound and MDA-modified proteins), pyroptosis activation proteins (NLRP3, cleaved caspase-1, N-terminal GSDMD) and cytotoxic cytokines (IL-1β, IL-6, TNF-α), but reversed the down-regulation of cytoprotective cytokine IL-10. Intranasal LiCl in RFV also alleviated the loss of the postsynaptic synapse protein PSD-95, but not synapsin-1, in aged 5XFAD mice. Blood level of the kidney function marker creatinine was significantly increased in 5XFAD than in WT mice in an age-dependent manner and this elevation was abolished by intranasal delivery of LiCl in RFV. Intranasal LiCl in RFV for 12 weeks in both WT or 5XFAD mice did not affect blood biomarkers for thyroid function, nor did it affect smell or muscle function or body weight.

**Conclusion:** Intranasal administration of LiCl in RFV significantly decreased lithium blood concentrations and increased brain/blood lithium concentration ratio, in comparison to its oral administration. Intranasal administration of LiCl in RFV robustly protected against both memory loss and depressive-like behavior, while had no side effects concerning thyroid and kidney toxicity in 5XFAD mice. These lithium-induced beneficial effects were strongly associated with lithium’s suppression of InsP_3_R-1 Ca^2+^ channel receptor increase, pathological neuroinflammation and activation of the pyroptosis pathway, as well as the loss of some synaptic proteins. Intranasal delivery of lithium salt in RFV could become an effective and potent inhibitor of pathological inflammation/pyroptosis in the CNS and serve as a new treatment for both AD-associated dementia and depression with minimal unwanted side effects including peripheral organ toxicity.

## Introduction

The importance of developing an effective treatment for Alzheimer’s disease (AD), a devastating neurodegenerative disease and a major public health problem affecting 6 million patients in USA alone[1], cannot be overemphasized. The major AD pathology is the progressive dementia, which is the predominant cause of dementia in aged population[1]. To be effective disease-modifying drugs for AD treatment, the drugs should be capable targeting critical upstream pathology[2] and associated majority of the 25 proposed AD pathologies[3]. In addition, the drugs must have increased therapeutic windows, and be well tolerated by AD patients without significant side effects/toxicity with chronic use[4].

Although many mechanisms have been proposed to cause neurodegeneration and synapse damage in AD, pathological inflammation and associated programmed cell death by pyroptosis has been considered one of the primary pathologies[5, 6]. In AD or many other neurodegenerative diseases, the concentration of the neurotransmitter glutamate in the synapse space can be as high as 100 mM[7], leading to over activation of N-methyl D-Aspartate (NMDA) receptor (NMDARs) overactivation. This receptor overactivation causes massive Ca^2+^ influx from extracellular space into cytosol, resulting in Ca^2+^ imbalance and excitotoxicity[8]. The Ca^2+^ dysregulation can be further aggravated by excessive Ca^2+^ release from the primary intracellular Ca^2+^ store endoplasmic reticulum (ER) via over activation of inositol trisphosphate receptors (InsP_3_ R)[9] and ryanodine receptors[10, 11]. This upstream Ca^2+^ dysregulation results in downstream mitochondria dysfunction[12], pathological reactive oxygen species (ROS) production and lipid peroxidation[13], leading to activation of pyroptosis regulatory proteins (NLR family pyrin domain containing 3 (NLRP3), cleaved caspase-1, N-terminal of Gasdermin D (GSDMD)), release of neurotoxic cytokines (IL-1β and IL-18) and programmed cell death by pyroptosis[14]. Pyroptosis plays a critical role in the pathophysiology of dementia[15] and associated depression behavior[16], and could be a therapeutic target[6] for both cognitive dysfunction and psychiatric disorders.

Lithium salt is a well-established first-line drug for the treatment and prophylaxis of mania and depression in bipolar disorder. Although neither the etiology of bipolar disorder nor lithium’s therapeutic mechanism is clear, numerous reports have shown that lithium has robust neuroprotective properties against a variety of preclinical models of neurodegenerative diseases[17], including AD[18]. Multiple mechanisms have been proposed for the efficacy of lithium in treating bipolar patients in clinics and cognitive impairments in experimental AD settings. These proposed mechanisms include the ability of lithium to ameliorate upstream Ca^2+^ dysregulation by inhibiting NMDAR[19] and suppressing InsP_3_ R[20]. Multiple downstream pathological events—including oxidative stress, inflammation, synapse loss and neuronal destruction by pyroptosis—are considered the primary mechanisms by which lithium may alleviate both cognitive dysfunction and depression behaviors in AD and other neurodegenerative diseases[21]. However, major concerns or limitations for lithium therapy in bipolar disorder patients is its narrow therapeutic window, proneness to side effects/toxicity, and intolerance in patients with chronic use[22, 23]. As lithium’s neuroprotection is typically dose-dependent, the effective dose for neuroprotection in the CNS will likely cause intolerable side effects and/or organ toxicity. Thus, a new strategy to promote its CNS therapeutic effects, while minimizing its peripheral side effects and organ toxicity is crucial to promote lithium use for treatment of both dementia and depressive disorder in AD or other neurodegenerative diseases.

Emerging evidence supports that intranasal application of molecules, compounds or even cells can enter the brain by using olfactory nerves, trigeminal nerves, and nasal lymphatics/vasculatures, bypassing the blood-brain barrier and circumventing most of the peripheral side effects[24]. Additionally, the use of nanoparticle has the potential of enhancing drug delivery to the brain via uptake by the brain endothelial cells to overcome the blood-brain barrier[25]. Our pioneer studies demonstrated dantrolene dissolved in RFV (Ryanodex), a form of crystalline nanoparticles [26], significantly increased dantrolene brain/blood concentration ratio[27, 28]. However, whether other chemicals, like lithium salt, dissolved in RFV will also increase drug brain/concentration ratio remained unknown. Therefore, we first compared the changes of lithium concentration in brain versus blood after administration of oral or intranasal LiCl in RFV or in water. We then investigated the beneficial versus side effects and the potential mechanisms by inhibition of inflammatory pyroptosis using intranasal treatment with LiCl in RFV to ameliorate both cognitive dysfunction and depression behavior in WT versus 5XFAD mice.

## Materials and Methods

### Animals

All the procedures were approved by the Institutional Animal Care and Use Committee (IACUC) of the University of Pennsylvania. Four pairs of 5XFAD mice (B6SJL-Tg (APPSwFlL on, PSEN1*M146L*L286V) 6799Vas/Mmjax) and wild type (WT) mice (B6SJLF1/J) were purchased from the Jackson Laboratory (Bar Harbor, ME) and bred. These 5XFAD transgenic mice over-expressed mutant human APP with the Swedish (K670N, M671L), Florida (I716V), and London (V717I) Familial Alzheimer’s Disease (FAD) mutations along with human PS1 harboring two FAD mutations, M146L and L286V. C57BL/6J mice were purchased from Jackson lab (Bar Harbor, Main, U.S.A.) and were used in studies of lithium brain/blood concentration ratio. Food and water were available in the animal cage. All mice were weaned no later than one month of age and were genetically identified by polymerase chain reaction (PCR) analysis before weaning. At this time, mice were divided into different cages according to age and gender, with no more than five mice per cage. Both male and female mice were used in this study.

### Lithium chloride preparation and administration

LiCl (Sigma, Aldrich, Co, St. Louis, MO, U.S.A.) was dissolved in Ryanodex Formulation Vehicle (RFV: 125 mg mannitol, 25 mg polysorbate 80, 4 mg povidone K12 in 5 ml of sterile water and pH adjusted to 10.3) as in our previous publications[27, 28], or in water. The dose of LiCl for both intranasal and oral administration was 3 mmol/kg. Intranasal administration of LiCl in RFV was performed following same protocol that we described previously[28]. The mouse was held and fixed by the scruff of their necks with one hand and with the other hand was given a total of 1 μl/g or 5 μl/g of body weight of LiCl in RFV or vehicle using a stock LiCl solution of 3 mol/ml and 0.6 mol/ml, respectively. The solution was slowly delivered directly into the mouse’s nose cavity. Care was taken to make sure that mice were minimally stressed, and that the solution stayed in the nasal cavity and did not enter the stomach or lung.

### Experimental treatment groups and drug administration Lithium pharmacokinetics studies

WT C57BL/6J mice at 2 months of age were divided into three groups: 1) intranasal administration of LiCl in RFV, 2). intranasal administration of LiCl in water, and 3) oral administration of LiCl in RFV. Brain versus plasma lithium concentrations were measured and brain/blood lithium concentration ratios were calculated at 30, 60 and 90 minutes after a single lithium administration, and results from different animal groups were compared.

### Measurements of lithium concentrations in the brain versus blood

The concentrations of lithium in the brain tissue and serum were measured by a lithium detection the Lithium Assay Kit (Colorimetric) (Abcam: Car# ab235613, Boston, U.S.A.) according to the manufacturer’s instructions. Briefly, brain tissues were homogenized in 0.5 N trifluoroacetic acid. Supernatant was collected after centrifugation at 10,000 x g for 5 minutes. The absorbance of all sample and standard curve wells was measured at both 540 nm and 630 nm in endpoint mode. The final lithium concentrations in the samples were calibrated and determined.

### Experimental lithium treatment groups

As demonstrated in Fig. 1, age-matched male and female mice were randomly divided into the experimental groups when they were genotyped around 1 month of age. The early treatment groups began treatment at 2 months of age and the late treatment group began at 9 months of age. The experimental lithium treatment groups included intranasal or oral LiCl in RFV and no treatment control. Mice were treated once per day and 5 times per week (Monday to Friday) for 12 weeks. LiCl in RFV was freshly prepared before each administration, as described in our previous studies[27, 28]. All behavioral tests were performed after completion of 12 weeks of treatment at 5 or 12 months of age for the early or late treatment groups respectively. Mice were euthanized at 6 or 13 months of age after 1 month of behavior tests, and the brain and blood were harvested to determine various biomarkers for the mechanistic study.

**Figure 1.**
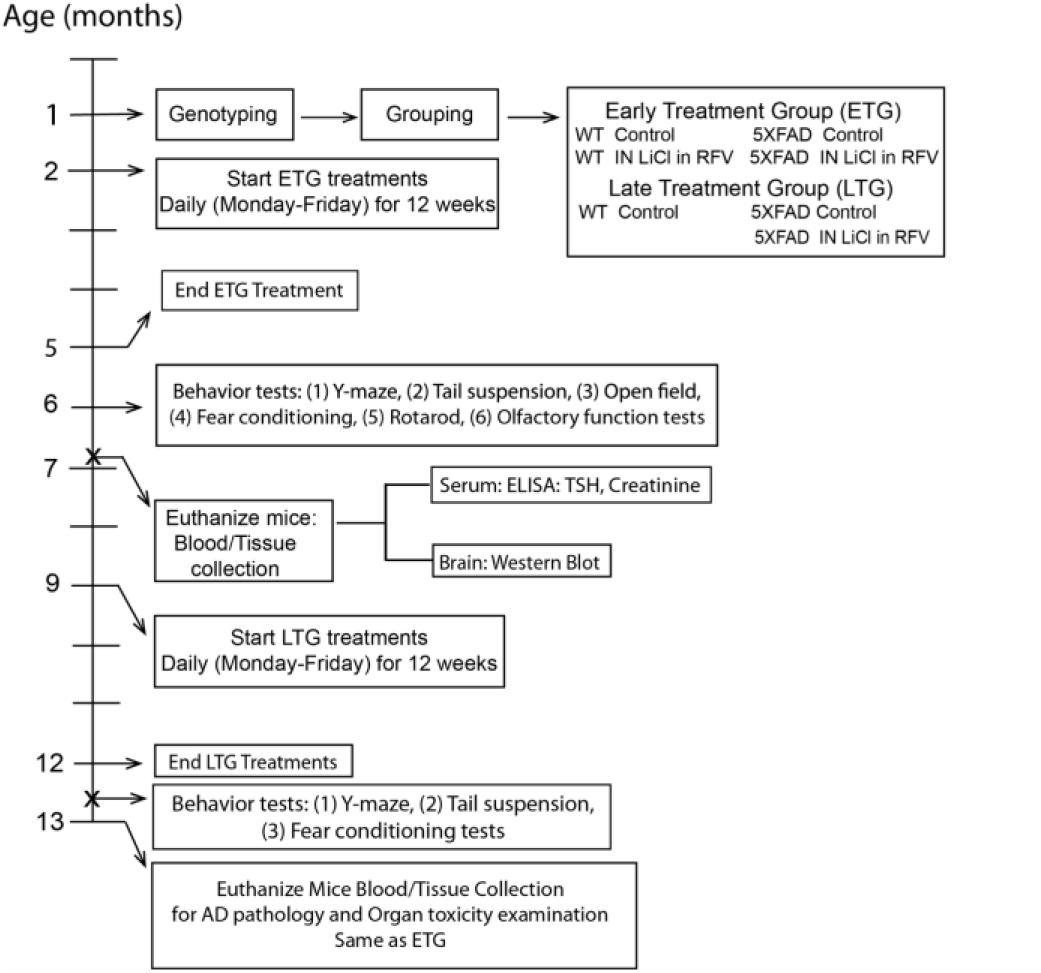
Experimental designs. LiCl: lithium chloride, RFV: Ryanodex Formulation Vehicle, ETG, early treatment group. LTG, late treatment group, ELISA: Enzyme linked immunosorbent assay, TSH; Thyroid Stimulating Hormone.

### Behavioral Assessments

#### Fear conditioning test

Memory was assessed at 6 or 13 months of age for the early or late treatment groups respectively. Both hippocampal-dependent and independent memory were assessed by the fear conditioning test, using the protocol we previously described[28]. Each day, animals were acclimated to the testing room at least 1 hour before the test. On the first day, each mouse was placed in the test chamber and went through three condition stimulation parings with a 60-second interval between each cycle. A 30-second tone of 2000 Hz and 85 dB was used as the tone stimulation and a 2-second electrical foot shock of 0.7mA was used as the shock stimulation. The mice were removed from the chamber 30 seconds after the last stimulation. On the second day, the contextual fear conditioning test was first performed to measure the hippocampal-dependent memory. The mouse was placed in the same chamber for 4 minutes with no tone or shock and then removed from the chamber. Two hours later, the cued fear conditioning test was performed to measure hippocampal-independent memory. The mouse was placed in another chamber that was different in size and smell using different cleaning solutions. There was no tone or shock during the first one minute. Later the mouse went through three cycles of the same tone with a 60-seconds interval between each cycle with the freezing time recorded. Animals were then removed from the chamber 60 seconds after the last tone. The ANY-maze controlled Fear Conditioning System consisted of a sound-attenuating chamber (Model: 46000-590, UGO Basile, Gemonio Italy) equipped with a video camera and ANY-maze software (V.4.99 Stoelting Co. Wood Dale, IL) which recorded the freezing time. The chamber was thoroughly cleaned between trials with a 75% alcohol solution on the first day in the training trials and on the second day in the contextual-fear conditioning test, and with water on the second day in the cued fear conditioning test. The investigator was blinded to the animal groups.

#### Y-maze test

Memory and learning were also assessed at 5- or 12-months age for the early or late treatment groups respectively. As described previously[29], memory was assessed using the Y-shaped maze with three grey, opaque plastic arms at a 120° angle from each other (Y-maze, NOLDUS Information tech Inc, Leesburg, VA, USA). Each day, animals were acclimated to the testing room at least one hour before the test. Each mouse was placed into the same arm of the maze and allowed to freely explore the three arms. The number of arm entries and the number of triads were recorded to calculate the percentage of alternation. The whole course was recorded by a camera and the results were calculated manually. The investigator was blinded to the animal groups.

#### Tail suspension test (TST)

Depression behavior was assessed 5 or 12 months of age for the early or late treatment groups respectively using tail suspension test described previously with modifications[30]. Briefly, we used specially manufactured tail suspension boxes made of carton with the dimensions (42 cm height X 18 cm width X 30 cm depth). To prevent animals from observing or interacting each other, each mouse was suspended within its own three-walled rectangular box. Each day, animals were acclimated to the testing room at least one hour before the test. Each mouse is suspended in the middle of the box and the width and depth were sufficiently sized so that the mouse cannot contact the walls. The bottom of each box had paper towel that collected feces or urine from the animals. We securely adhered both the mouse’s tail and the suspension bar to the box so that it was strong enough to carry the weight of the mouse being evaluated. A video camera was placed in position. After suspending the mouse on the suspension bar, the TST session was recorded. The total duration of the test was 6 minutes. After all sessions were finished, the videos were analyzed manually. During the behavioral analysis, the time that each mouse spent as mobile was measured and was subtracted from 360 seconds as the immobility time. The immobile time reflects the level of depression behavior. The investigator was blinded to the animal groups.

#### Rotarod test

Motor function was examined for muscle weakness and inco-ordination. The amount of time spent on the accelerating rotarod (IITC Series 8, Life Sciences, Woodland Hills, CA) was assessed for mice in the early treatment group at 6 months of age, as previously described[28]. Briefly, animals were acclimated to the testing room at least 1 hour before the testing. Two 60 second-training trials at a constant speed (9 rpm) were performed with a 30-minute interval. Then, three 120 second-testing trials were conducted at a gradually increasing speed (4–40 rpm) with a 60-minute interval between trials. The latency to fall from the rotarod was recorded automatically and analyzed.

#### Food buried test

Olfaction was assessed in the early treatment group at 5 months of age as previously described with modifications[27, 28]. On the first day, cookies were buried in the bedding of the home cages for 24 hours, and then the number of cookies consumed was recorded. Water was freely available during this time. The buried food test was conducted on the third day from 9–11 am. Mice were individually placed into a clean cage containing clean bedding with one cookie buried beneath the bedding in a corner. The latency for the animal to find the cookie (identified as catching the cookie with its front paws) was recorded manually. If the animal failed to find the cookie within 15 min, it would be placed back into its home cage. A clean cage and bedding were used for each animal and investigators were blinded to the experimental conditions.

#### Euthanasia and tissue collection

Mice of the early or late treatment groups were euthanized at 6 or 13 months of age respectively after all the behavior tests were finished. As we described previously[28], animals were deeply anesthetized with 2–4% isoflurane delivered through a nose cone, and the concentration was adjusted according to the animals’ response to a toe pinch. The animal’s skin was prepped, and an incision made to open the chest and expose the heart. Blood was collected for the serum study from the heart using a syringe equipped with a 27G needle, without heparin. The blood was centrifuged at 1,400 rpm at 4°C for 30 minutes; the supernatant was collected and frozen at –80°C. The animals were euthanized by trans-cardia perfusion and exsanguination with cold phosphate-buffered saline. The thyroid gland, kidney, and the brain were dissected. The brain was stored in –80°C for western blotting analysis.

#### Immunoblotting of proteins

Total brain tissues were extracted by homogenization using cold RIPA buffer (#9806S, Cell Technology, USA) supplemented with protease inhibitor cocktails (P8340 Roche). The brain homogenates were rocked at 4°C for 90 minutes and then centrifuged at 14,000 rpm (Brushless Microcentrifuge, Denville 260D) for 20 minutes at 4°C to remove cell debris. After collecting the supernatant, the protein concentration was measured using a BCA protein assay kit (Pierce, Rockford, IL 61101 USA). Briefly, equal amounts of protein (50 µg/lane) were loaded onto 4-20% gel electrophoresis of mini-protein TGX precast (Cat. #4561094, BIO-RAD) and transferred to polyvinylidene difluoride (PVDF) membranes (Immobilon-P, MERK Millipore, Ireland) using wet electrotransfer system (BIO-RAD, USA). Following the transfer membrane blocking with 5% BSA (Sigma-Aldrich) for 1 hour, the PVDF membrane was incubated overnight at 4°C with primary antibodies including InsP3R1, MDA- and 4-HNE-modified protein, NLRP3, Human procaspase-1/cleaved caspase-1 p20, GSDMD, cleaved GSDMD, IL-1β, IL-18, IL-6, TNF-α, PSD95, and GAPDH (Antibody details including their dilutions are shown in Table 1). Afterward, the membranes were incubated with secondary antibodies, including HRP-linked anti-mouse IgG1 and anti-rabbit IgG that was conjugated to horseradish peroxidase, then washed with Tris-buffered saline containing 0.2% Tween-20. After incubation with secondary antibodies, proteins band were visualized using ECL Western Blotting Detection Reagents (Cytiva, amersham, UK) and quantified for band intensity using ImageJ software (National Institutes of Health, Bethesda, MD, USA).

**Table 1:** Antibodies applied in immunoblotting.

| **Primary antibody** | **Sources** | **Dilutions** |
| --- | --- | --- |
| Anti- InsP3R1 | # PA1-901, Invitrogen | 1:1000 |
| Anti-MDA-modified protein | MA5-27560, Invitrogen | 1:1000 |
| Anti-4 Hydroxynonenal | (ab46545), Abcam | 1:1000 |
| Anti- NLRP3 | 68102-1-Ig; Proteintech | 1:2000 |
| Anti-Human procaspase-1/ cleaved caspase-1 p20 | #2225S, Cell signaling Technology, USA | 1:1000 |
| Anti-GSDMD | ab155233; Abcam | 1:1000 |
| Anti- Cleaved GSDMD | #36425S; Cell signaling Technology, USA | 1:1000 |
| Anti-IL-1β | 31202S, Cell signaling Technology, USA | 1:1000 |
| Anti-IL-18 | 10663-1-AP; Proteintech | 1:3000 |
| Anti-IL-6 | # M620; Invitrogen | 1:1000 |
| Anti-TNF-α | 60291-1-Ig; Proteintech | 1:2000 |
| Anti-GFAP | #3670; Cell signaling Technology, USA | 1:1000 |
| Anti-Iba1 | # 17198; Cell signaling Technology, USA | 1:1000 |
| Anti-IL-10 | #12163S, Cell signaling Technology, USA | 1:1000 |
| Anti-PSD95 | #2507; Cell signaling Technology, USA | 1:1000 |
| Anti-Synapsin-1 | #5297; Cell signaling Technology, USA | 1:1000 |
| GAPDH | MA5-15738, Thermo Fisher Scientific | 1:10,000 |
| **Secondary antibody** |  |  |
| Anti-Mouse IgG1; HRP -linked | # PA1-74421, Thermo Fisher Scientific | 1:10,000 |
| Anti-rabbit IgG, HRP-linked | #7074; Cell signaling Technology, USA | 1:1000 |

#### Serum Thyroid Stimulating Hormone (TSH) measurement

Serum TSH, a hormone whose level reflects the function of thyroid gland, was measured using a TSH Colorimetric Assay Kit (CHM01J578, Thomas scientific, Inc, Swedesboro, NJ, USA), an ELISA assay, according to the manufacturer’s instructions. We measured the serum thyroid stimulating hormone for the 5XFAD mice in the early treatment groups and all mice in the late treatments’ groups. Briefly, an aliquot of 10 μl serum was diluted with 40 μl of sample diluent and incubated for 30 minutes at 37°C. After repeated washings, 50 μl of HRP-conjugate reagent was added followed by 30-minute incubation at 37°C. The reaction was stopped with a stop solution and the optical density (OD) at 450 nm was measured within 15 minutes. A TSH standard curve was generated and the sample TSH concentration was calculated by using the linear regression equation of the standard curve.

#### Serum creatinine measurement

Serum creatinine, an indicator of kidney function, was measured using a Creatinine Colorimetric Assay Kit (ab65340, Abcam, Inc, Waltham, MA, USA) according to the manufacturer’s instructions. We measured the serum creatinine for the 5XFAD mice in the early treatment groups and all mice in the late treatment groups. Briefly, an aliquot of 50 μl serum was diluted in a 100 μl reaction mixture, including 42 μl Creatinine Assay Buffer, 4 μl Creatinase, 2 μl Cr Enzyme Mix, and 2 μl Creatinine Probe., The OD at 570 nm was measured at 60 minutes after incubating the reaction at 37°C. A creatinine standard curve was generated and the sample creatinine concentration was measured in a linear range of the standard curve. The trendline equation standard curve (Sa) was calculated from the standard curve data. The creatinine concentration was calculated using the formulation: creatinine concentration =Sa/50 nmol/μl.

#### Statistics

All data was represented as mean ± standard errors mean (SEM). Statistical analyses were employed with GraphPad Prism (Version 9.3.1, CA, USA). Comparisons of more than two groups were conducted by two-way ANOVA with Tukey’s multiple comparison test. P-value of <0.05 was considered statistically significant.

## Results

### Intranasal delivery of LiCl in RFV (Ryanodex Formulation Vehicle) significantly increases the brain/blood lithium concentration ratio

Wild-type (WT) mice at 2 months of age were treated with LiCl (3 mmol/kg) dissolved in RFV by intranasal application. For comparisons, we are also intranasally treated these mice with LiCl dissolved in water or orally with LiCl in RFV. Lithium concentrations in the brain and serum were measured and brain/blood concentration ratios were calculated. The lithium brain concentrations in mice treated with intranasal LiCl in RFV were higher than those treated with intranasal LiCl in water or oral administration of LiCl in RFV, only at 2 hours, but not 30 or 60 minutes after treatment (Fig. 2A). The lithium blood concentrations at three time points after intranasal LiCl in RFV administration were significantly lower than those observed after oral lithium LiCl in RFV administration (Fig. 2B). At 2 hours, the brain/blood lithium concentration ratio in the group of LiCl in RFV was significantly higher than that of the oral LiCl in RFV or intranasal LiCl in water group. (Fig. 2C).

**Figure 2.**
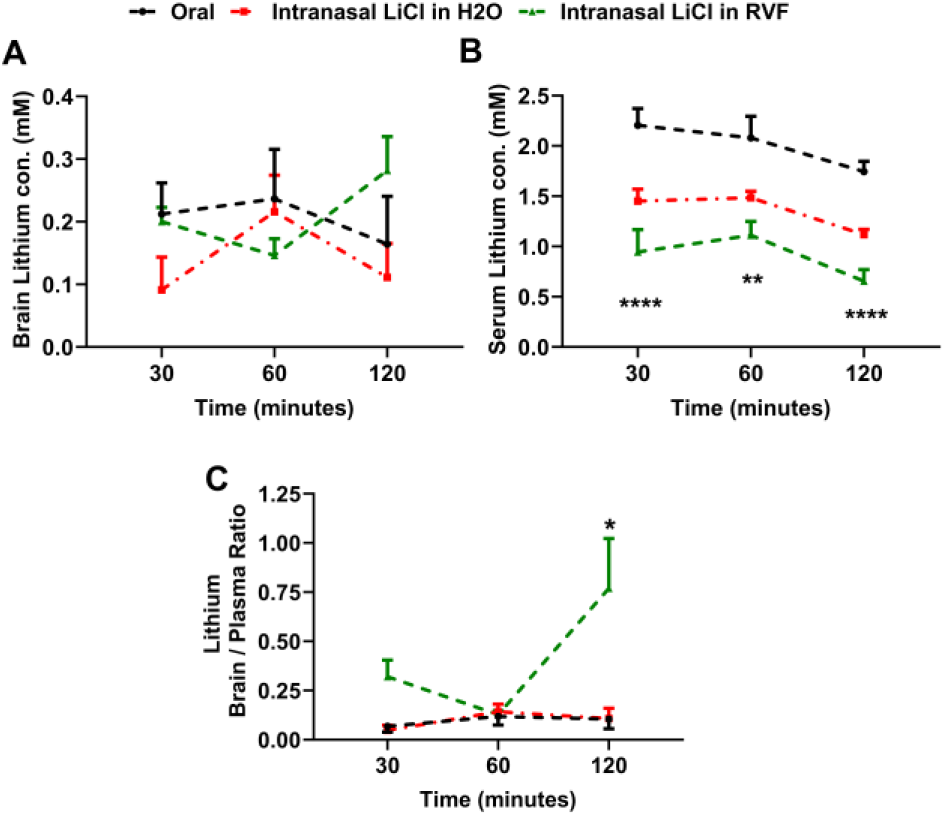
Intranasal LiCl in RFV increased the brain/blood concentration ratio compared to oral LiCl in RFV in adult wild type mice. WT mice at 2 months of age were given LiCl (3 mmol/kg) dissolved in Ryanodex formulation vehicle (RFV) intranasally (green), orally (black), or intranasally but dissolved in water (red). Lithium concentrations in the brain (**A**) and serum (**B**) were measured and brain/blood concentration ratios were calculated (**C**), at 30, 60 and 120 minutes after drug administration. Data are Means±SEM (N=6) and analyzed by Two-way ANOVA followed by Tukey’s multiple comparison test for statistical significance. **P<0.01, ****P<0.001, compared to oral administration of LiCl in RFV (black) in Fig. 2B. *P<0.05 compared to oral administration of LiCl in RFV (black) and intranasal administration of LiCl in water (green) in Fig. 2C.

### Intranasal LiCl in RFV alleviates both memory loss and depression behavior in both young adult and aged 5xFAD mice

Next, we accessed the effects of intranasal delivery of lithium in RFV on behaviors of 5XFAD mice. The freezing time in fear condition test reflects the level of memory; the longer the freezing time means better memory. The contextual fear conditioning (CFC; hippocampus-dependent) test demonstrated significant memory loss in young 6 months of age 5XFAD, which could be abolished by 12 weeks of intranasal LiCl in RFV treatment (Fig. 3A). In aged mice at 12 months of age in the late treatment group, the hippocampus-independent FC-cued test indicated a significant memory loss in aged mice and could be fully corrected by intranasal LiCl in RFV treatment for 12 weeks in aged mice (Fig. 3B). However, the cognitive function determined by the Y-maze did not detect a memory loss, nor did the intranasal or oral LiCl in RFV affect cognitive function in 5XFAD mice at either 6 (Supplemental Fig. 1A) or 12 (Supplemental Fig. 1B) months of age.

**Figure 3.**
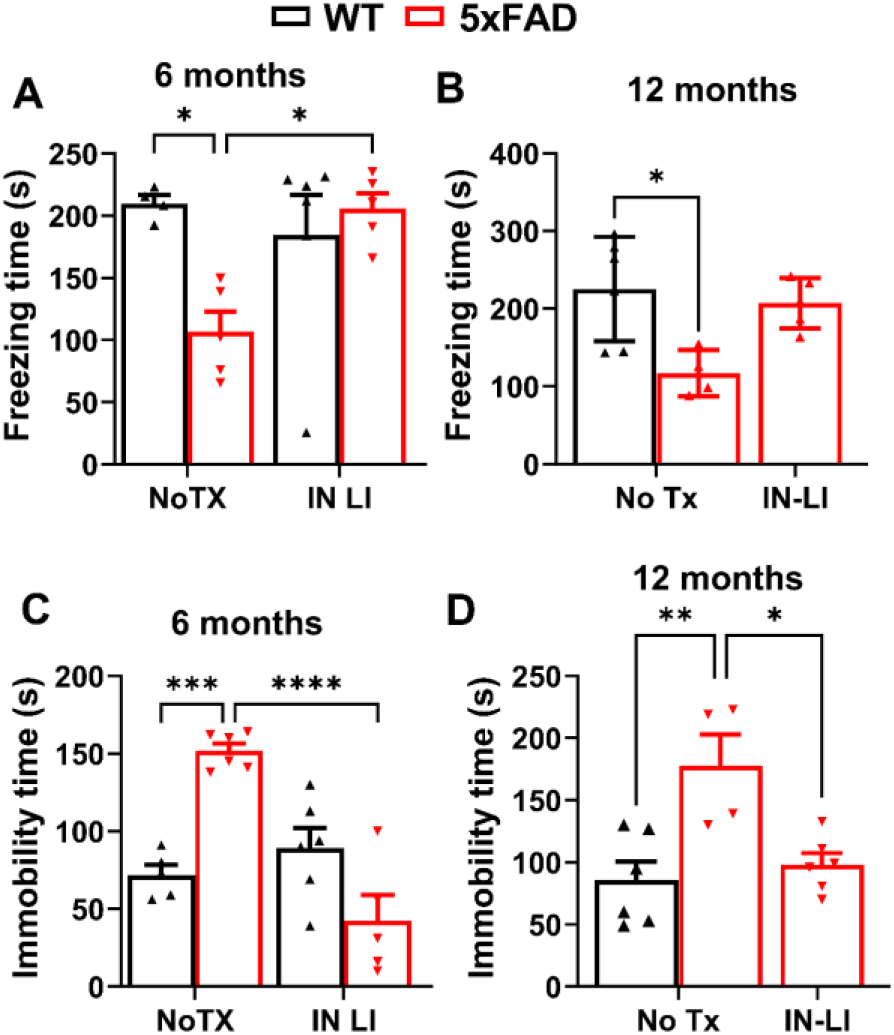
Intranasal LiCl in RFV significantly alleviated memory loss and depressive behavior in 5XFAD mice. WT or 5XFAD mice at 2 or 9 months of age received either no treatment (No TX) or intranasal LiCl in RFV (IN LI, 3 mmol/kg), daily, Monday to Friday, for 12 consecutive weeks. Cognitive function was determined by fear conditioning at 6 (**A** hippocampus-dependent) or 12 (**B**, hippocampus-independent) months of age. Depressive behavior was determined by tail suspension test at 6 **(C)** or 12 **(D)** months of age. Data represents Means±SEM from 4-6 mice and were analyzed using the 2-way ANOVA followed by Tukey’s multiple comparison test **(A, B, C, D)**. * P<0.05, ** P<0.01, *** P<0.001, **** P<0.0001.

In the tail suspension test (TST) to measure levels of depression behavior, immobility time represented the level of depression behavior; the more immobility time means higher level of depressive behavior. The immobility times in 5XFAD mice were significantly higher than in WT control mice in both young adult (Fig. 3C) and aged groups (Fig. 3D). Notably, intranasal LiCl in RFV treatment for 12 consecutive weeks abolished the increased immobility time in both young (Fig. 3C) and old (Fig. 3D) 5XFAD mice, returning it to the levels of WT mice. Overall, these results demonstrate that intranasal delivery of LiCl in RFV can abolish both memory loss and depression-like behavior in both young and old 5XFAD mice.

### Intranasal Lithium in RFV inhibits the pathological increase of type 1 InsP_3_ R (InsP_3_ -1) in aged 5XFAD mice

As an important AD pathology, InsP_3_R-1 activity is significantly increased in AD cells[9] or animal models[31]. Therefore, we determined the effects of intranasal LiCl in RFV treatment on changes of InsP_3_ -1 protein levels using immunoblotting. Intranasal LiCl in RFV treatments for 12 weeks abolished the pathological increase of InsP_3_ -1 protein in aged 5XFAD mice brains (Fig. 4A, B). These results suggest that intranasal delivery of lithium salt in RFV nanoparticles blocks the pathological elevation of InsP_3_R-1 protein, the critical Ca^2+^ channel that leads to excessive Ca^2+^ release from the ER. This action in turn inhibits an upstream critical Ca^2+^ dysregulation core pathology and ultimately ameliorates both memory loss and depression behavior in these AD mice.

**Figure 4:**
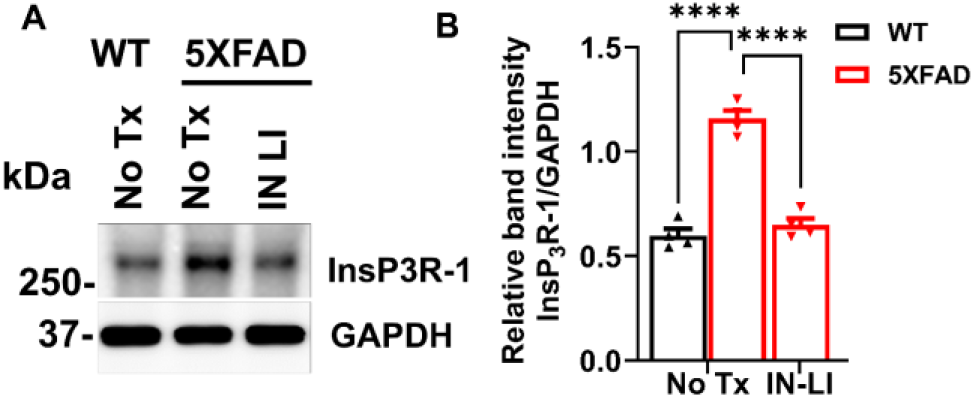
Intranasal LiCl in RFV inhibited the pathological increase of type 1 InsP_3_ R (InsP_3_R-1) proteins in the brain of old 5XFAD mice. WT or 5XFAD mice at 9 months of age received either no treatment (No Tx) or intranasal LiCl in RFV (INLI, 3 mmol/kg) daily for 12 consecutive weeks. Mice brains harvested at 13 months of age for immunoblotting of the expression level of InsP_3_ Rs-1 (**A**) and statistical analysis of relative band intensity of InsP_3_ Rs-1/GAPDH (loading control). (**B**). Data are Means±SEM from 4 separate mice in each treatment group (N=4) and were analyzed using the two-way ANOVA followed by Tukey’s multiple comparison test. ****P<0.0001.

### Intranasal lithium in RFV suppresses the oxidative stress in 5XFAD mice

It has been reported that oxidized biomolecule by-products, produced in the neuronal cells due to the overproduction of malondialdehyde (MDA)-modified proteins, lead to increased ROS in AD brains[32]. Protein-bound 4-hydroxynonenal (4-HNE) is also a widely used oxidative stress marker[33]. Therefore, we measured protein oxidation by determining oxidized biomolecule by-products, MDA-modified proteins, and the protein bound 4-HNE. Both protein-bound 4-HNE and MDA-modified proteins were significantly increased in 5XFAD mice, in comparison to the WT control. Intranasal LiCl in RFV abolished the pathologically increased protein-bound 4-HNE (Fig. 5A, B) and MDA-modified proteins (Fig. 5C, D).

**Figure 5.**
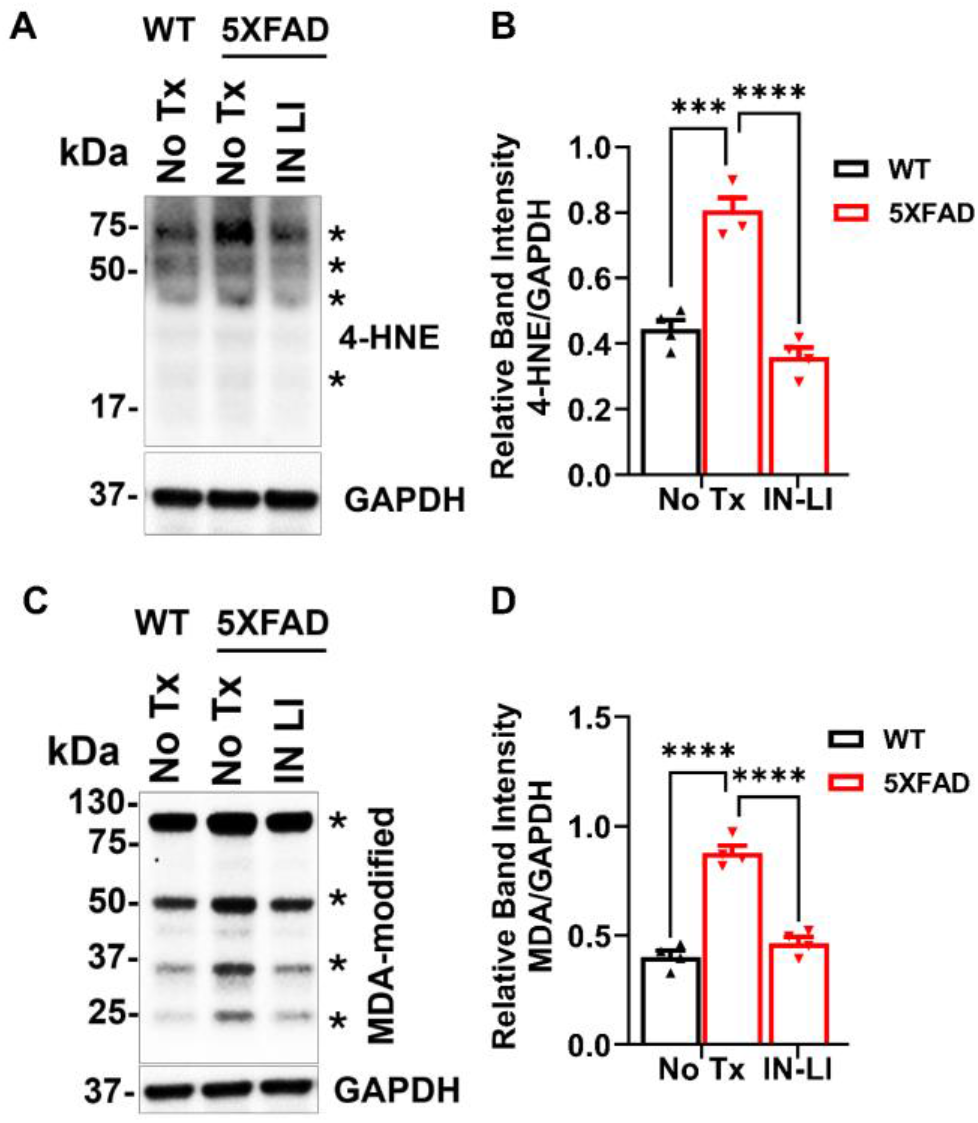
Intranasal LiCl in RFV blocked oxidative stress in the brain of old 5XFAD mice. WT or 5XFAD mice at 9 months of age were received either no treatment (No Tx) or intranasal LiCl in RFV (INLI, 3 mmol/kg) for 12 weeks. Mice brains were harvested at 13 months of age for immunoblotting of the lipid peroxidation product 4-HNE- (**A, B**) or MDA- (**C, D**) modified proteins, and combined of 4-HNE- or MDA-modified proteins indicated by asterisks (**\*)** are quantified for statistical analysis. Data indicates Means±SEM from 4 separate mice in each treatment group (N=4) and were analyzed using the two-way ANOVA followed by Tukey’s multiple comparison test. ****P<0.0001.

### Intranasal lithium in RFV inhibits pathological activation of pyroptosis pathway in the brain of 5XFAD

We measured the protein levels of primary regulators of pyroptosis, including NLR family pyrin domain containing 3 (NLRP3), caspase-1, Gasdermin D full-length (GSDMD-FT) and N-terminal Gasdermin D (N-GSDMD) in old adult WT versus 5XFAD mice. The core inflammasome complex protein NLRP3 was significantly increased in the brain of 13-month-old 5XFAD mice compared with the WT control, and this increase was reversed by the intranasal treatment with LiCl in RFV for 12 consecutive weeks (Fig. 6 A, D). Intranasal LiCl in RFV treatment robustly inhibited pathological elevation of cleaved caspase-1 (Fig. 6B, E), had no effects on level of GSDMD-full length (Fig. 6C, F), but inhibited elevation of N-terminal Gasdermin D (N-GSDMD, Fig. 6G, J), which is the core on plasma membrane to release neurotoxic cytokines (IL-1β and IL-18) from cytosol into extracellular space[5, 34]. Consistently, intranasal LiCl in RFV robustly inhibited the pathologically increased IL-1β (Fig. 6H, K) and IL-18 (Fig. 6I, L) in 5XFAD mice brains, two critical cytokines contributing to activation of pyroptosis[14]. Overall, these results suggest that intranasal LiCl in RFV significantly suppresses the activation of the pyroptosis pathway in the brain of aged 5XFAD mice.

**Figure 6.**
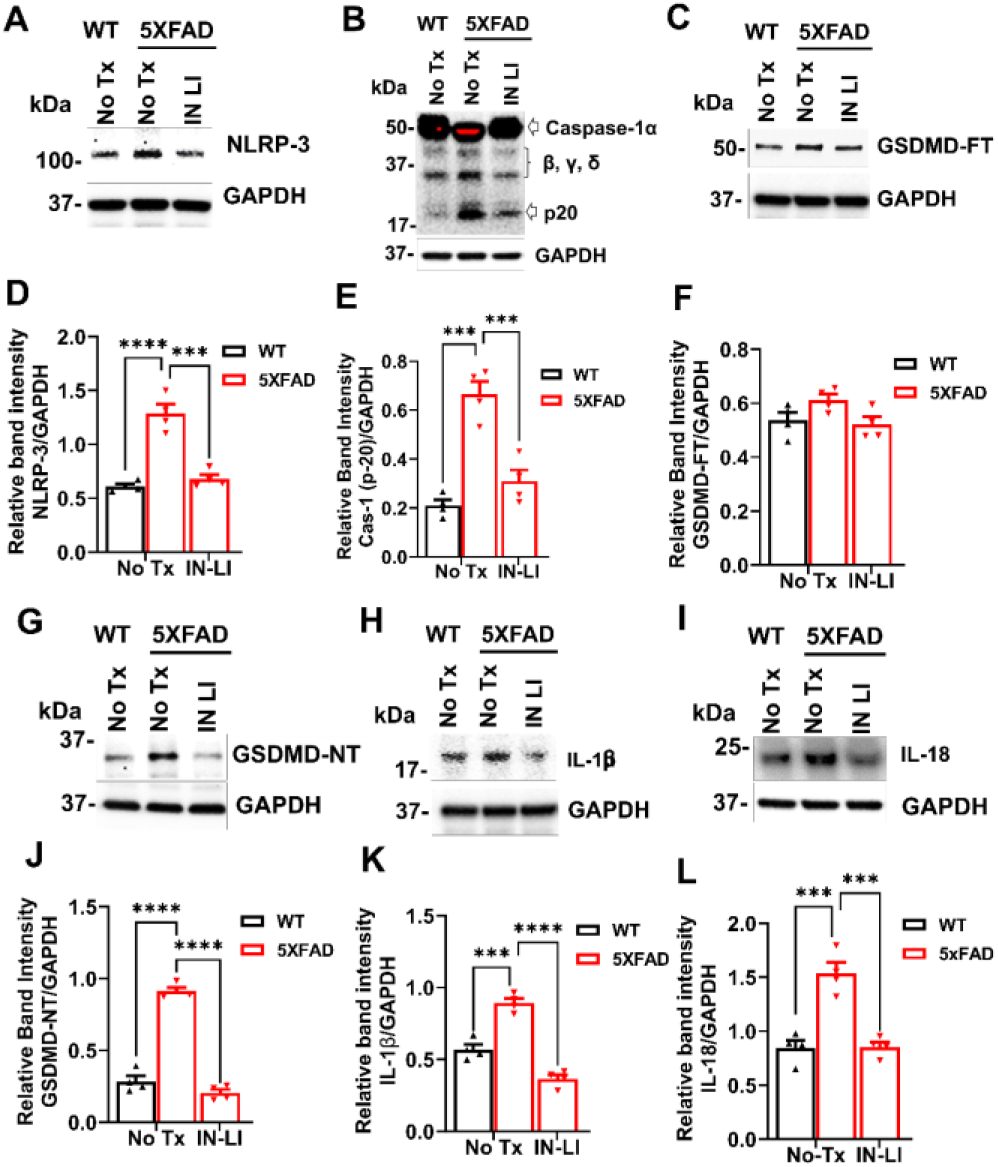
Intranasal LiCl in RFV abolished pathological activation of pyroptosis in 5XFAD mice brains. WT or 5XFAD mice at 9 months of age either no treatment (No Tx) or intranasal LiCl in RFV (INLI, 3 mmol/kg), daily for 12 consecutive weeks. Mice brains harvested at 13 months of age. Changes in key regulatory proteins of the pyroptosis pathway were identified. NLRP3: NLR family pyrin domain containing 3 **(A, D)**, Caspase-1 (p-20) **(B, E)**, GSDMD-FL: full length GSDMD Gasdermin D **(C, F)**, GSDMD-NT: N terminal Gasdermin D **(G, J)**, IL-1β **(H, K)**, and IL-18 **(I, L)** of representative Western blots and related statistical analysis. Data are Means±SEM from 4 separate mice brains (N=4) and were analyzed using two-way ANOVA for each protein followed by Tukey’s multiple comparison test. ***P<0.001, ****P<0.0001.

### Intranasal lithium in RFV alleviates the pathological elevation of neurotoxic cytokines and inhibited astrogliosis and microgliosis

Both astrogliosis and microgliosis contribute to pathological inflammation[35, 36]. GFAP, a biomarker protein for astrogliosis, was significantly increased in 5xFAD mouse brain compared to WT controls, but this increase was robustly inhibited by intranasal treatment with LiCl in RFV (Fig. 7A, C). Similarly, the biomarker protein for microgliosis, IBA-1, was significantly elevated in the brain of 5XFAD mice compared to the WT control and this increase was also abolished by intranasal LiCl in RFV (Fig. 7 B, D). Pathological elevation of cytotoxic cytokines plays a critical role on programmed death by pyroptosis[14]. We have determined changes of critical neurotoxic (IL-6, TNF-α) and neuroprotective (IL-10) cytokines in the brain of aged 5XFAD mice, both with and without intranasal LiCl in RFV treatments. Cytotoxic cytokines (TNF-α, IL-6) were pathologically elevated (Fig.8A, D, B, E), while the cytoprotective cytokine (IL-10) was markedly decreased (Fig. 8C, F). These imbalances were corrected by intranasal LiCl in RFV treatment. Altogether, our results indicate that intranasal treatment with lithium ion in RFV suppresses the pathological neuronal inflammation in aged 5XFAD mice.

**Figure 7.**
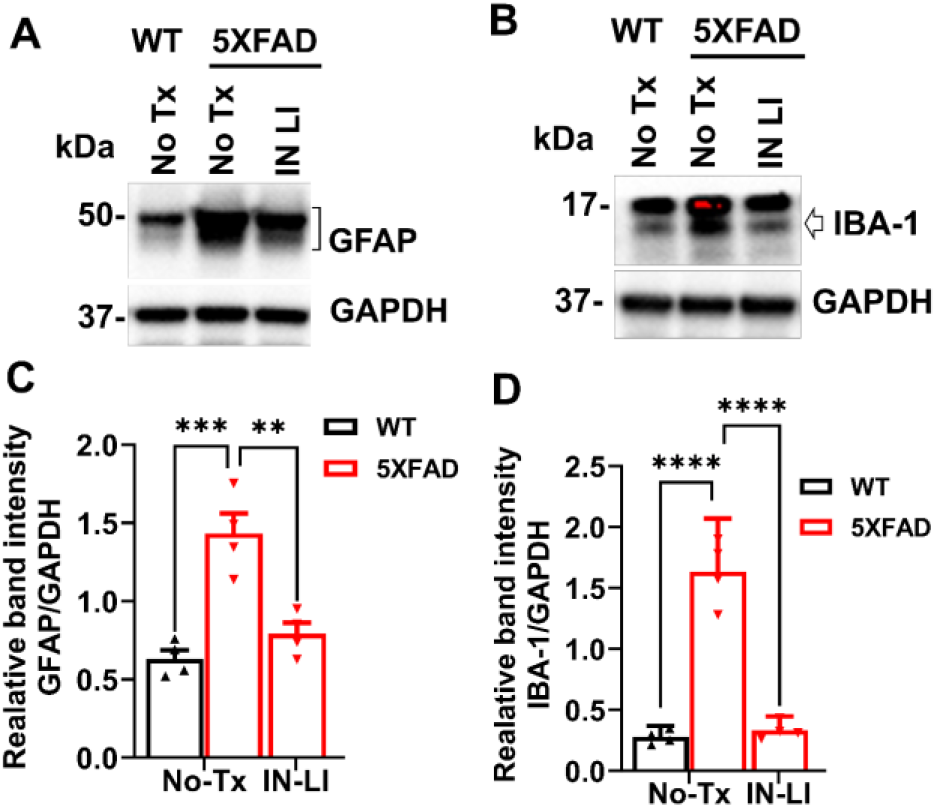
Intranasal LiCl in RFV suppressed astrogliosis/microgliosis in the 5XFAD mice brains. WT or 5XFAD mice at 9 months of age received either no treatment (No Tx) or intranasal LiCl in RFV (INLI, 3 mmol/kg) for 12 consecutive weeks. Mice brains were harvested at 13 months for immunoblotting to determine levels of protein biomarkers of astrogliosis (GFAP) **(A, C)** and microgliosis (IBA-1) indicated by arrow **(B, D)**. Data are Means±SEM from 4 separate mice brains (N=4) and analyzed using two-way ANOVA followed by Tukey’s multiple comparison test (C, D). **P<0.01, ***P<0.001, ****P<0.0001.

**Figure 8.**
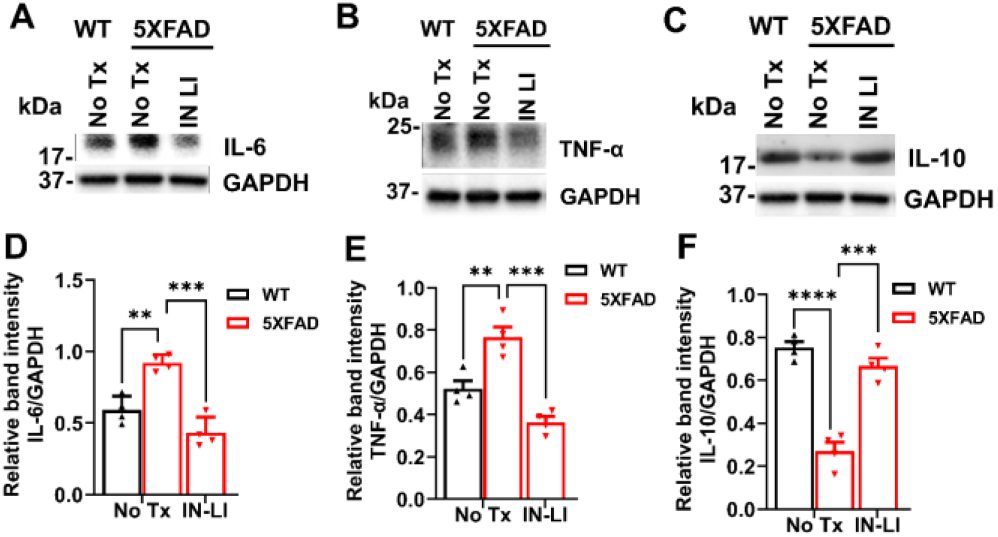
Intranasal LiCl in RFV normalized the upregulation of neurotoxic cytokines and downregulation of neuroprotective cytokine in 5XFAD mice brains. WT or 5XFAD mice at 9 months of age received either no treatment (No Tx) or intranasal LiCl in RFV (INLI, 3 mmol/kg) for 12 consecutive weeks. Mice brains were harvested at 13 months of age for immunoblotting of protein biomarkers of neurotoxic IL-6 **(A, D)**, TNF-α **(B, E)** and neuroprotective IL-10 **(C, F)** cytokines. Data are Means±SEM from 4 separate mice brains (N=4) and analyzed using two-way ANOVA followed by Tukey’s multiple comparison test (**D, E, F**). ** P<0.01, ***P<0.001, **** P<0.0001.

### Intranasal lithium in RFV prevents the loss of some postsynaptic synapse protein in the brain

Normal synapse structure plays an important role in synapse function, while their destruction contributes to cognitive dysfunction and depression behavior[37]. Levels of the postsynaptic synapse protein PSD-95 were significantly decreased in 13-month-old, aged 5XFAD mice compared to WT controls. This decrease was reversed by intranasal treatment with LiCl in RFV for 12 weeks from 9 to 12 months of age (Fig. 9A, C). However, intranasal lithium treatment did not inhibit the loss of synaptin-1 protein loss (Fig. 9B, D)

**Figure 9:**
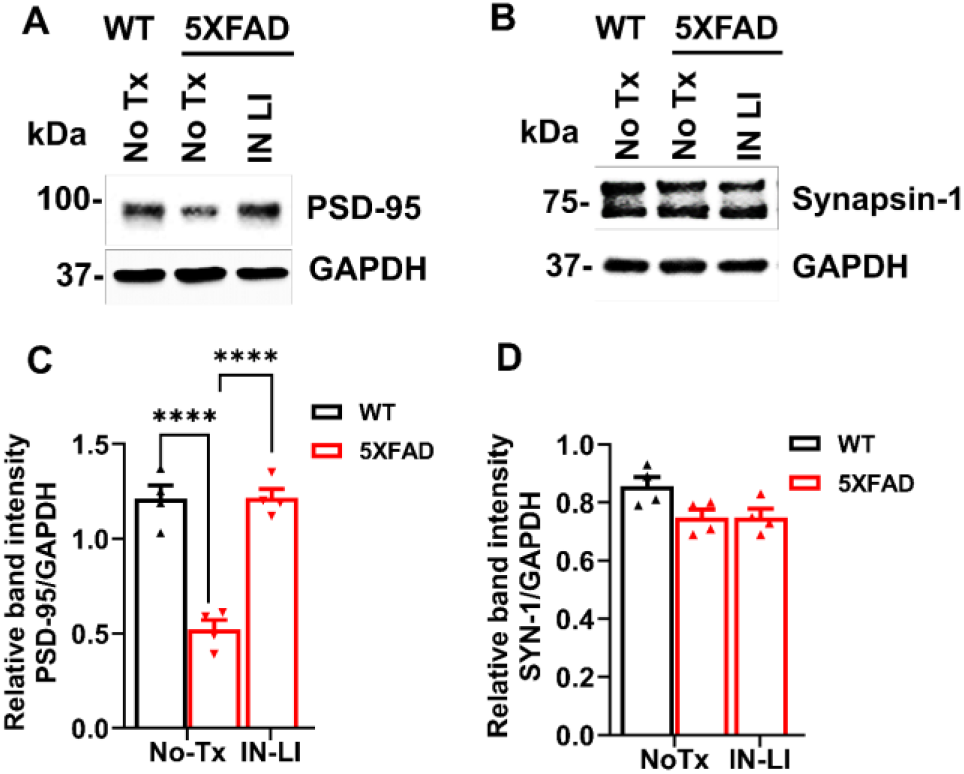
Intranasal LiCl in RFV restored the loss of synapse proteins in 5XFAD mice brains. WT or 5XFAD mice at 9 months of age received either no treatment (No Tx) or intranasal LiCl in RFV (INLI, 3 mmol/kg), daily for 12 consecutive weeks. Mice brains were harvested at 13 months for immunoblotting of PSD-95 **(A, C)** and Synapsin-1 **(B, D)**. Data are Means±SEM from 4 separate mice brains (N=4) and analyzed using two-way ANOVA followed by Tukey’s multiple comparison test. **** P<0.0001 (C, D).

### Chronic administration of intranasal lithium in RFV did not cause thyroid or kidney toxicity or side effects

Chronic use of lithium, especially at a high dose, is prone to cause dysfunction of peripheral organs, notably thyroid and kidney[38]. We therefore used ELISA calorimetric kits to measure the biomarkers reflecting the function of thyroid and kidney. Intranasal LiCl in RFV treatment for 12 consecutive weeks in either the early treatment group (from 2 to 5 months of age) or the late treatment group (from 9 to 12 months of age), did not cause significant changes in levels of the blood biomarkers for thyroid (Fig. 10A, B), but blocked kidney dysfunctions detected in both young adult (6 months of age) and aged (13 months of age) 5XFAD mice (Fig. 10C, D). Additionally, intranasal LiCl in RFV for 12 weeks in the early treatment groups did not cause significant changes of motor or nose smell function determined by the rotarod (Supplemental Fig. 2A) and food buried test (Supplemental Fig. 2B), respectively. No significant changes in body weight were detected during intranasal treatment with LiCl in RFV in the late treatment group (Supplemental Fig. 2C, D). These results suggest that chronic intranasal LiCl in RFV treatment is safe without significant organ side effects or organ toxicity.

**Figure 10.**
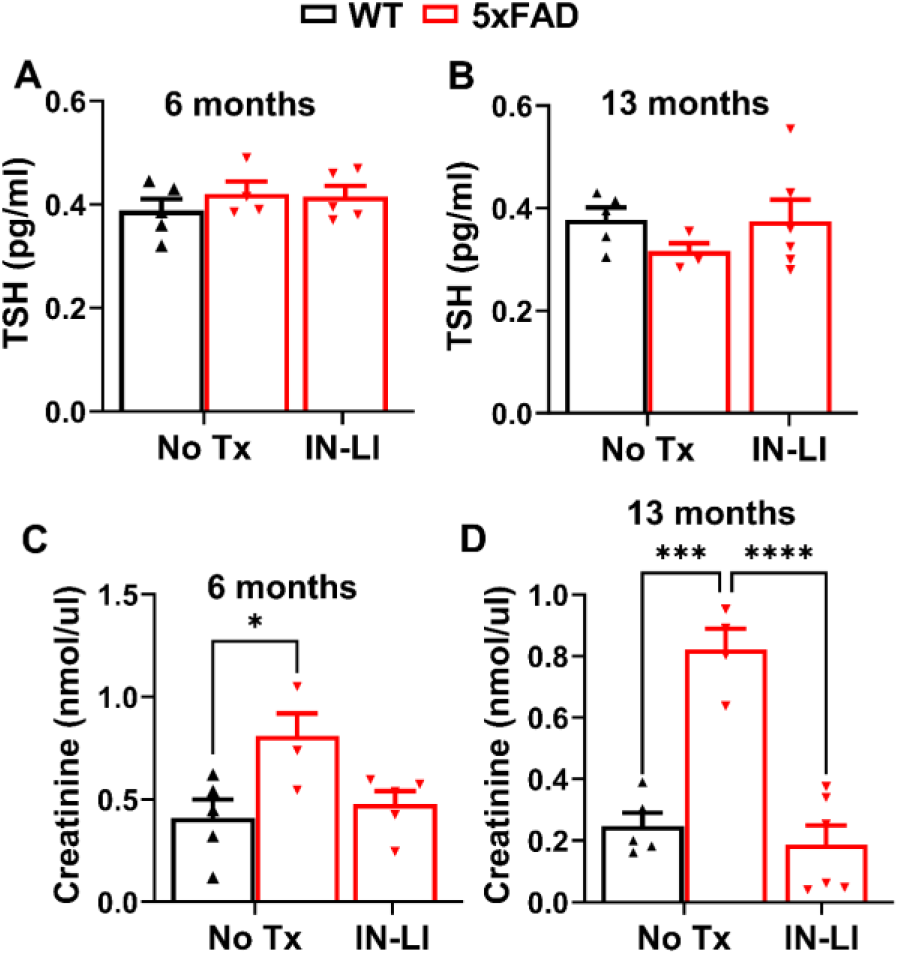
Intranasal LiCl in RFV did not affect thyroid function and protected against kidney dysfunction in 5XFAD mice. WT or 5XFAD mice at 2 or 9 months of age received either no treatment (No Tx) or intranasal LiCl in RFV (INLI 3 mmol/kg) for 12 consecutive weeks. Mice bloods were then harvested at the age of 6 (**A, C**) and 13 (**B, D**) months of age. Respective blood biomarkers for the function of thyroid (**A, B**) (TSH: thyroid stimulating hormone) and kidney (**C, D**) (creatinine) were determined by ELISA at the indicated ages. Data are Means±SEM from blood of 4-6 separate mice (N=4-6) and analyzed using two-way ANOVA followed by Tukey’s multiple comparison test. *P<0.05, ***P<0.001, **** P<0.0001.

## Discussion

This study employed a novel approach using intranasal delivery of lithium chloride in RFV to demonstrate a marked increase in lithium brain/blood concentration ratio of lithium ion, compared with results using either intranasal lithium chloride in water or oral lithium chloride in RFV. Intranasal chronic treatment with lithium in RFV robustly blocked the memory loss detected using the FC test and depression-like behavior using the tail suspension test in young and old 5XFAD mice. Intranasal chronic lithium chloride in RFV also effectively protected against the pathological increase of InsP_3_ R Ca^2+^ channel protein levels, oxidative stress, pathological inflammation, and programmed cell death by pyroptosis. Importantly, these beneficial effects in 5XFAD mice were associated with no toxic effects on thyroid function and protective effects on kidney function after chronic use.

Although lithium has been demonstrated to have some neuroprotective efficacy in some preclinical AD cell or animal models and even in some AD patients[18, 39], its notorious narrow therapeutic window and being prone to induce side effects or organ toxicity may limit its potential future utility for treating of AD patients, especially under the need of chronic use[23]. Intranasal administration of drugs promotes their penetration into the brain, limits the entrance to the peripheral blood system, increases the duration on the nasal surface, and maintains a high level of concentrations in the CNS[40]. Also, intranasal administration of drugs in nanoparticle formulation further promotes penetration into the CNS and has been increasingly used for treatment of CNS disorders[41]. We have previously demonstrated that dantrolene, an ER Ca^2+^ release regulator, dissolved in RFV, a formulation of crystalline nanoparticles[26], significantly increased the drug delivery to the brain[27, 28], and attenuated memory loss in 5XFAD mice without serious adverse side effects following protracted treatment [28]. In the current study, our novel approach of using intranasal administration of LiCl in RFV significantly increased the lithium brain/blood concentration ratio two hours after administration, compared to either intranasal LiCl in water or oral LiCl in RFV. It is not clear if lithium dissolved in RFV also form nanoparticles like Ryanodex, which needs further studies. The particle structure of LiCl or other forms of lithium salt (lithium carbonate, lithium orotate etc.) dissolved in RFV should be further studied to optimize its brain/blood lithium concentration ratio for a strong therapeutic efficacy in the CNS with minimal side effects or organ toxicity in the peripheral system. Nevertheless, a higher brain/blood lithium concentration ratio indicates a promotion of lithium’s therapeutic effects in the CNS, with a lower risk of peripheral system side effects or organ toxicity. Therefore, a lower dose of lithium in RFV via intranasal administration can be used to achieve same therapeutic effects rather than by the commonly used oral lithium administration in RFV or water. It is important to study whether RFV will also assist other drugs difficult to pass BBB to penetrate brain more efficiently.

Although dementia is the primary symptom in AD patients, the high incidence of depression disorder may deteriorate dementia and even form a vicious cycle to worsen each other. Thus, a drug capable of effectively treating both dementia and depression behavior would promote high therapy efficacy in AD patients. Lithium is a first-line drug to treat mania and depression in bipolar disorder, but its use to treat depression in AD is less well studied. The efficacy of lithium to treat cognitive dysfunction is not consistently found in AD patients[17]. Considering both the necessity for prolonged treatment duration and the narrow therapeutic window of lithium, the chronic use of high-dose lithium for treatment of AD may face limitations. With the novel approach of intranasal lithium in RFV administration, this study demonstrates a robust protection against both cognitive dysfunction and depression behavior. Given the need of chronic treatments of most neurodegenerative diseases, especially AD, achieving higher brain/blood concentration ratios through intranasal lithium chloride administration compared to the oral approach provides a better opportunity to minimize the lithium dose. This approach aims to keep the dose as low as possible, reducing the risk of thyroid and kidney toxicity and other side effects with chronic lithium administration. The therapeutic efficacy and benefit of intranasal administration of various lithium formulations in RFV should be further studied in different AD animal models and eventually evaluated in AD patients, as well as in patients with psychiatric disorders, such as bipolar disorder.

Accumulating studies suggest that inflammation plays a significant role in AD pathology and contributes to both cognitive dysfunction and depression behavior. Overactivation of NMDAR on plasma membranes and/or RyRs on ER membranes in AD collectively contributes to the upper stream pathological increase in cytosolic and mitochondrial Ca^2+^ concentrations. This increase results in downstream mitochondrial dysfunction, oxidative stress, and subsequent activation of the NLRP3 inflammasome complex and caspase-1. This cascade also leads to an elevation of cytokines IL-1β and IL-18, cleavage of GSDMD into an ion channel N-terminal to release cytotoxic cytokines, and ultimately programmed cell death by pyroptosis[5]. Although lithium has been demonstrated to be neuroprotective by inhibition of inflammation, its effects and mechanisms on inflammation associated pyroptosis has not been well investigated. This study demonstrates that intranasal LiCl nanoparticles significantly inhibits upstream Ca^2+^ dysregulation by ameliorating the pathological increase of InsP_3_R-1 and abolishes downstream oxidative stress, pathological inflammation, pyroptosis, and synapse destruction in 5XFAD mouse brains. In consistence, intranasal LiCl in RFV may effectively treat both memory loss and depression behavior by its potent inhibition of inflammatory pyroptosis and synapse proteins loss.

Calcium dysregulation, along with associated inflammation and pyroptosis, also plays critical role in pathologies of neurodegenerative diseases, such as stroke, Parkinson disease, Huntington disease, seizure, and Amyotrophic Lateral Sclerosis (ALS) [5, 42]. These neurodegenerative diseases often are chronic diseases and need prolonged treatments. The novel approach involving intranasal lithium nanoparticle administration is expected to promote its therapeutic effects in the CNS while minimizing peripheral side effects and organ toxicity, especially under prolonged treatment for various brain disorders[23]. Thus, we propose that intranasal LiCl in RFV may provide neuroprotection not only in AD, but other commonly seen neurodegenerative diseases and psychiatric disorders.

Our study is the first to demonstrate that blood creatinine, a kidney function biomarker, was significantly increased in 5XFAD mice compared to the WT control, especially in aged 5XFAD mice, indicating a kidney dysfunction. Blood creatinine levels are associated with the blood biomarkers for neurodegeneration, including neurofilament light chain(NFL)[43], amyloid protein[44], and phosphorylated tau[43], as well as the risk of dementia[45]. An important finding in this study is that both intranasal and oral lithium in RFV, especially via the intranasal approach, abolished the abnormally increased blood creatinine level, suggesting that lithium can protect kidney dysfunction in AD, and function as another mechanism to ameliorate dementia. As GSDMD, a critical pyroptosis player, and inflammation play important role contributing to kidney damage and dysfunction[46], the potent inhibition of pyroptosis and inflammation by lithium may explain, at least in part, its protective effects on kidney function in this study. Notably, kidney dysfunction is one of the organ toxicities associated with chronic and high-dose lithium treatment for bipolar disorder, and the blood lithium concentration in this study is low. It is important to promote the use of low doses of lithium for chronic treatment of AD, neurodegenerative diseases, or psychiatric disorders to minimize side effects and organ toxicity. Intranasal lithium nanoparticles, with their potential to reduce side effects and organ toxicity compared to commonly used oral administration, warrant further investigation.

It should be noted that there is inconsistency on effects of lithium on cognitive function determined by fear conditioning versus Y-Maze tests. Only fear conditioning test but not Y-Maze test demonstrated the inhibitory effects of lithium on cognitive dysfunction in 5XFAD mice. A plausible reason for this inconsistency is the sensitivity of behavioral tests to determine cognitive function is quite different. Our previous study demonstrated a significant memory loss determined using fear conditioning test but not by Morris Water Maze test in 5XFAD mice[28]. Also, there is a significantly memory loss in triple transgenic AD mice at young adult age[47], but not aged mice[48], determined by same Morris Water Maze. Nevertheless, the result in this study is consistent with our previous study[28] that there are significant memory loss in old adult triple transgenic AD mice and 5XFAD mice determined by fear conditioning. These results suggested that fear conditioning test is sensitive to detect cognitive dysfunction in old adult or aged mice and should be elected and the results should be confirmed together with other sensitive behavioral tests.

This study has the following limitations: 1). The sample size in each experimental group is small. However, lithium demonstrated significantly statistically therapeutic effectiveness, suggesting its robust potency of neuroprotection in this AD animal model. 2). We were unable to measure the changes of cytosol versus mitochondria Ca^2+^ levels in the brain tissue, but with lithium’s blockade of the increase of InsP_3_ Rs protein levels, the study indirectly suggests lithium inhibits upstream Ca^2+^ dysregulation. 3). We did not measure the contents of reactive oxygen species (ROS) concentrations in brains directly but used the changes of oxidized lipid membrane proteins as indirect indication of oxidative stress. 4). We only examined the inflammation biomarkers and pyroptosis pathway activation proteins in old, but not young-adult brains. As 5XFAD mice at 6 months of age already developed full AD pathologies and cognitive dysfunction, effectiveness of lithium to ameliorate memory loss and depression behavior suggest that intranasal lithium chloride nanoparticles can be considered a disease-modifying drug in AD. We predict that the lithium will have similar neuroprotective effects against inflammation and pyroptosis and associated synapse destruction in young-adult 5XFAD mice. Additionally, we did not administer intranasal lithium nanoparticle vehicle treatment in older adult wild type mice because the treatment did not significantly alter memory or depression behavior in young-adult WT mice.

In conclusion, this study strongly suggests that chronic intranasal LiCl in RFV ameliorates both cognitive dysfunction and depression behavior in 5XFAD mice, without affecting thyroid, smell, and motor function, and while also protecting kidney function. Lithium neuroprotection in 5XFAD mice was associated with its robust inhibition of pathologically increased InsP_3_ Rs protein levels, oxidative stress on lipid membrane proteins, pathological inflammation, cell death by pyroptosis and synaptic protein loss. This study will inspire future research work investigating neuroprotective versus side effects of intranasal lithium salt in RFV in AD and other neurodegenerative diseases or psychiatric disorders.

## Acknowledgements

The research was performed in the lab of Dr. Huafeng Wei and should be attributed to the Department of Anesthesiology, University of Pennsylvania. We appreciate technical support from Yutong Yi from the Department of Anesthesiology and Critical Care.

## Author contributions

H.W. conceived and designed the study. P.B, W.Z, G.L, B.J, R.V, R.C, K.K, L.S.L, Y.W, J.L, and H.W. conducted experiments and acquired the data, P.B, W.Z, G.L, B.J, R.V, R.C, K.K, L.S.L, and H.W. analyzed data. P.B, W.Z, G.L, R.C, K.K, L.S., D.-M.C, and H.W contributed to the manuscript preparation. All the authors reviewed and approved the final manuscript.

## Data Availability

Data is available upon request.

## Supplemental Figures and Legends

**Supplemental Figure 1.**
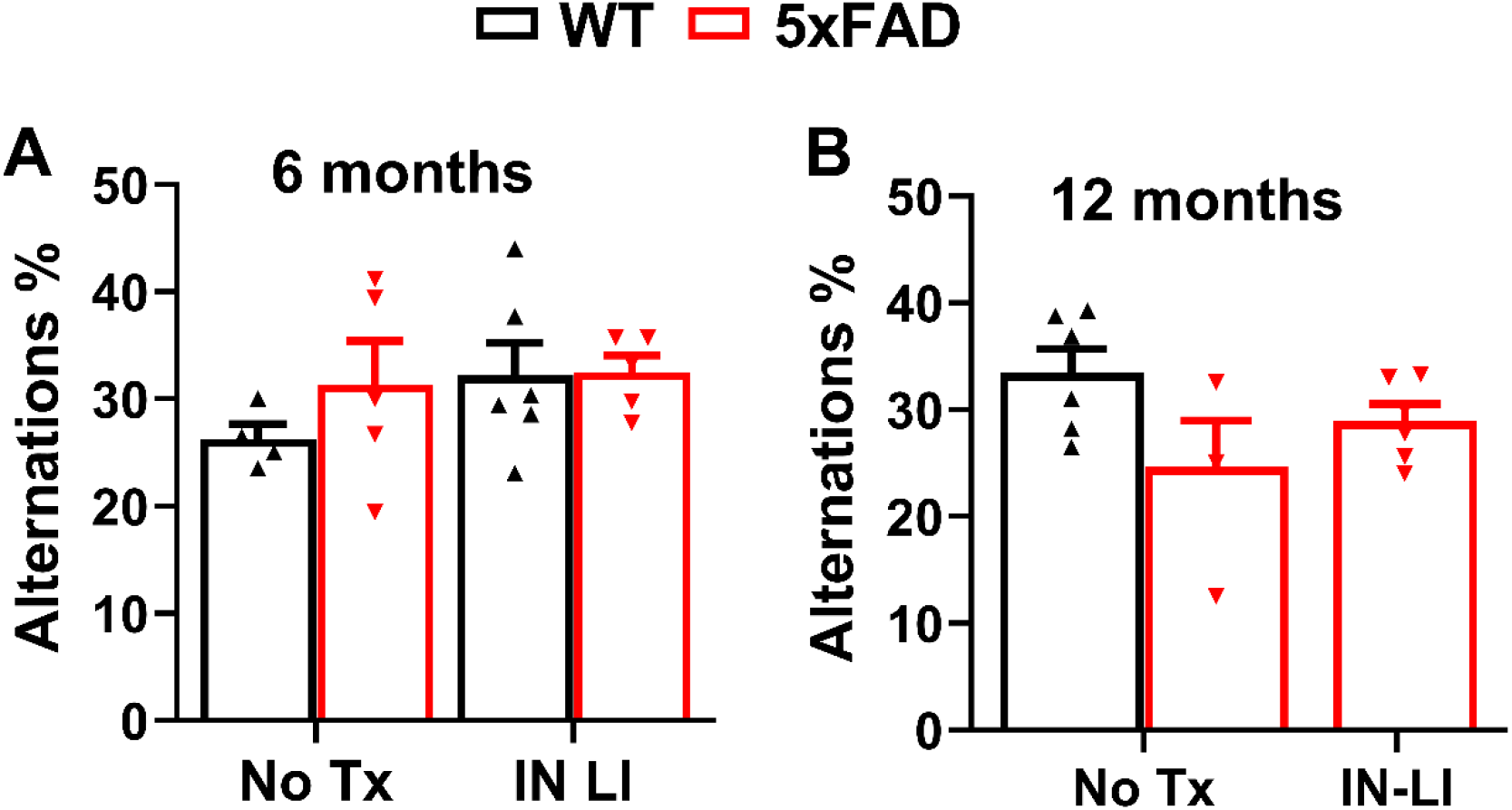
Effects of intranasal LiCl in RFV on cognitive function assessed by Y-maze test in 5XFAD mice. WT or 5XFAD mice at 2 or 9 months of age received either no treatment (No Tx) or intranasal LiCl (IN LI, 3 mmol/kg) dissolved in RFV daily, Monday to Friday, for 12 consecutive weeks. Y-maze tests were performed at 6 (**A**) and 12 months of age (**B**), after completion of treatments. Data represents Means±SME from 4-6 mice (N=4-6) and was analyzed using the 2-way ANOVA followed by Tukey’s multiple comparison test.

**Supplemental Figure 2:**
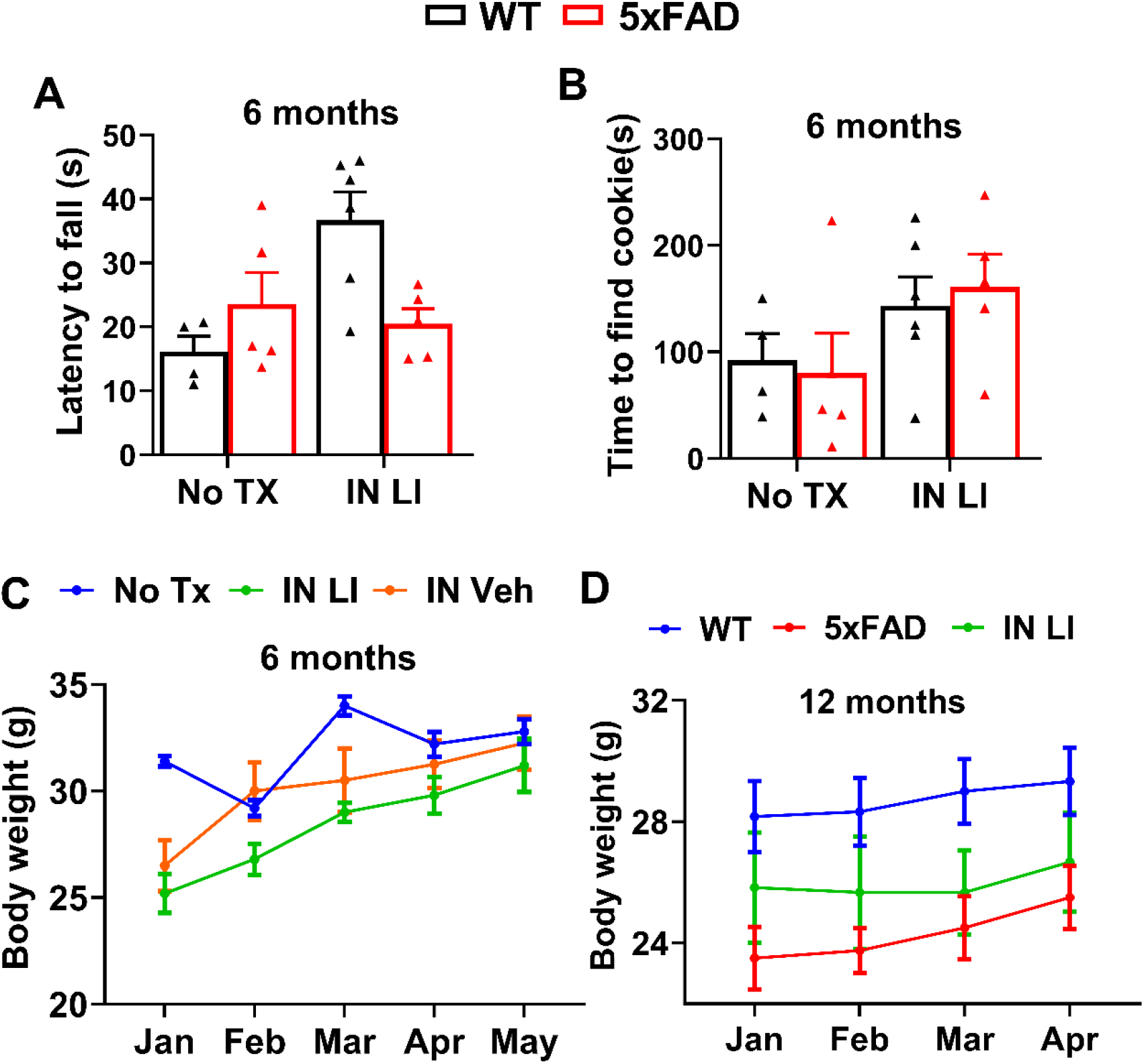
Effects of intranasal LiCl in RFV on motor and smell functions and body weight. WT or 5XFAD mice received either no treatment (No Tx) or intranasal LiCl in RFV (INLI, 3 mmol/kg) for 12 consecutive weeks from 2 to 5 months of age. Motor (**A**) and olfactory (**B**) functions were determined by rotarod test and food buried test, respectively at 6 months of age. Body weights were determined monthly from the initiation of lithium treatment at 2 months of age **(C)** until the end of all behavioral tests at 12 months of age(**D**). Data are Means±SEM from 4-6 mice in each group.

